# Hypoxia modulates human pulmonary arterial adventitial fibroblast phenotype through HIF-1α activation

**DOI:** 10.1101/2025.01.27.635152

**Authors:** Jennifer L Philip, Christine A Caneba, Laura R Caggiano, Nandhini Prakash, Tik-Chee Cheng, Katherine A. Barlow, Tasneem Mustafa, Diana M Tabima, Timothy A Hacker, Kristyn S Masters, Naomi C Chesler

## Abstract

Hypoxic pulmonary hypertension (HPH) develops in association with diseases characterized by low oxygen levels leading to pulmonary artery (PA) narrowing and death. Hypoxia has been linked to increased PA collagen and changes in PA adventitial fibroblast (PAAF) metabolism. However, the mechanisms by which hypoxia regulates PAAF function are unknown. Hypoxia-inducible factor-1α (HIF-1α) is a subunit of a transcription factor that is degraded in normoxia but stabilized in hypoxia and is involved in extracellular matrix remodeling by fibroblasts. We examined the role of hypoxia and HIF-1α in regulating PAAF function. Human PAAF (HPAAF) were cultured in normoxic and hypoxic conditions. Cells were further treated with HIF1-α inhibitor or no drug. Protein expression, mRNA expression, enzyme activity, and metabolite concentration were examined. Male C57BL6/J mice were exposed to 0 or 10 days of hypoxia after which right ventricular hemodynamics and tissue metabolism were assessed. Hypoxia led to an increase in collagen content and decrease in matrix metalloproteinase-2 (MMP2) activity. HIF-1α inhibition limited collagen accumulation and restored MMP2 activity. HPAAF demonstrated elevated lactic acid concentration and decreased ATP in hypoxia. HIF-1α inhibition blunted these effects. Mice exposed to hypoxia developed significant elevation in right ventricle systolic pressures and had decreased ATP levels in pulmonary tissue. This study investigated the mechanisms by which hypoxia drives HPAAF-mediated collagen accumulation and metabolic changes. We identify the key role of HIF-1α in regulating changes. These findings provide important insights into understanding HPAAF-mediated PA remodeling and help identify possible novel therapeutic targets.

## Introduction

Pulmonary hypertension is a rare but deadly disease, the hallmark of which is narrowing of the pulmonary arteries that can trigger right heart failure and mortality. Hypoxic pulmonary hypertension (HPH), or hypertension caused by low oxygen conditions, develops in the setting of many other diseases, such as obstructive sleep apnea (1), cystic fibrosis (2), and chronic obstructive pulmonary disorder (COPD) (3). The presence of pulmonary hypertension (PH) in these diseases is associated with increased mortality (4). Therefore, it is important to understand the processes involved in the development of hypoxic pulmonary hypertension in order to develop treatments to halt disease progression.

An important consequence of PH is pulmonary arterial stiffening, which increases right ventricular (RV) afterload and contributes to RV failure (5–7). HPH in mice leads to collagen accumulation throughout the pulmonary arterial wall, which correlates with pulmonary arterial stiffening (6, 8). In neonatal calves, HPH leads to collagen accumulation in the pulmonary artery adventitia (9). Hypoxia has been shown to activate adventitial fibroblasts, leading to an increase in extracellular matrix components and the accumulation of collagen in the adventitia (10–12). Excess collagen accumulation can occur due to increased collagen production or decreased collagen degradation. In HPH, the specific mechanism by which hypoxia leads to collagen accumulation in the pulmonary artery remains unclear.

Pulmonary arterial adventitial fibroblasts (PAAF) are the primary source of arterial collagen. Until recently, pulmonary arterial remodeling in pulmonary hypertension, including collagen accumulation, was believed to result from endothelial cell (EC) mechanotransduction of stress and subsequent signaling to medial smooth muscle cells (SMC) and PAAF, known as the “inside-out” mechanism (11). However, newer studies support an “outside-in” mechanism, in which mechanotransduction by PAAF and transmission to medial SMCs and intimal ECs leads to remodeling throughout the vessel wall (11, 13). Based on these recent findings, this paper focuses on the “outside-in” mechanism of pulmonary arterial remodeling in HPH.

Hypoxia-inducible factor-1 (HIF-1) is a heterodimeric transcription factor involved in transcribing genes related to proliferation, remodeling, and metabolism (14–16). HIF-1α is a subunit of this transcription factor that is normally degraded in normoxia, but stabilized in hypoxia (15). HIF-1α activation and/or expression is linked to changes in extracellular matrix remodeling in fibroblasts (14) and plays a role in cardiovascular disease (17) and pulmonary hypertension (18, 19). Also, HIF-1α activation and/or expression alters the metabolism of cancer cells (20, 21), which are also responsible for large amounts of collagen remodeling in hypoxia (22, 23). Specifically, increased activation or expression of HIF-1α led to increases in glycolytic rate in cancer cells (20, 21, 24).

Here, we investigate the direct effects of hypoxia on PAAF collagen production and degradation and how these are linked to changes in fibroblast metabolism. In addition, we explore the role of HIF-1α in hypoxia-induced increases in collagen synthesis by PAAF and alterations in PAAF metabolism. Finally, we seek confirmation of in vitro results in vivo.

## Materials and Methods

### Cell Culture

Human pulmonary artery adventitia fibroblasts (HPAAF) were purchased from ScienCell (Carlsbad, CA) and maintained in fibroblast growth medium (ScienCell) containing 2% fetal bovine serum, 1% fibroblast growth supplement, 20 U/mL penicillin, and 20 μg/mL streptomycin. Cells of passage 3 were used for all experiments. Cells were incubated in hypoxia using StemCell Technologies (Vancouver, Canada) hypoxia chamber flushed with gas containing 1% oxygen, 5% carbon dioxide, and 94% nitrogen, according to the manufacturer’s instructions.

### RT-qPCR

HPAAF were seeded at a density of 60,000 cells/mL in a 24-well plate in serum-free media and incubated overnight to allow cell attachment. Media was then changed to serum-free media or serum-free media containing 30 µM HIF-1α inhibitor (CAS 934593–90-5, Santa Cruz, Dallas, TX). Cells were placed in either a standard incubator (normoxia) or in the hypoxia chamber (hypoxia) 48 hours. Cells were then washed with PBS and RNA isolation was performed using RNeasy Mini Kit (Qiagen, Valencia, CA). RNA was reverse transcribed into cDNA using a High Capacity cDNA Reverse Transcription Kit (Applied Biosystems, Foster City, CA) as previously described (25). Real-time PCR was performed using TaqMan Gene Expression Master Mix and TaqMan assay primers (Collagen 1: Hs00164004_m1, GAPDH: Hs02786624_g1) according to the manufacturer’s instructions (Invitrogen Life Technologies, Carlsbad, CA). All PCRs were performed using the Applied Biosystems StepOne Plus Real-Time PCR System (Foster City, CA). Changes in mRNA expression were determined by the comparative threshold cycle method (26). Data were normalized to GAPDH and are expressed as fold change with respect to the normoxia control

### MMP Detection

HPAAF were seeded at a density of 60,000 cells/well in a 24-well plate in serum free media and cultured in the same manner as in the PCR experiment. After 48 hours, media was collected for immediate quantification of MMP activity, and cell lysate was collected for measurement of relative amounts of MMP2 via Western blot. MMP activity was measured using an MMP fluorogenic assay (Abcam, Cambridge, UK). Briefly, MMPs were activated by incubating collected media with 2mM 4-aminophenylmercuric acetate (APMA) for three hours. 50 µL of substrate was added to 50 µL of APMA-activated media in a 96-well plate. After 1 hour of incubation, fluorescence was read at 485 nm excitation/527 nm emission wavelengths using a Fluoroskan Ascent Microplate Fluorometer (Thermo Labsystems, Philadelphia, PA). Background fluorescence was subtracted based on control media samples.

Standard procedures were used to prepare samples for MMP2 analysis via Western blot. Briefly, cells were washed once with PBS and lysed with 63.3% glycerol, 2% SDS, 50 mM Tris-HCl (pH 6.8), 10 μg/mL aprotinin, 10 μg/mL leupeptin, 1 μg/mL pepstatin, 1 mM PMSF, 50 U/mL Benzonase nuclease, 1X Phosphatase Inhibitor III, and 5X Phosphatase Inhibitor II (Boston BioProducts, Ashland, MA). Lysates were vortexed, placed on ice for 30 min, and centrifuged at 21,000 x g for 15 min at 4°C. Equal volumes of lysates were separated via electrophoresis using NuPage® Novex® 4-12% Bis-Tris Protein Gels (Life Technologies) and transferred onto nitrocellulose membranes (Bio-Rad). Immunoblotting was performed using the Odyssey Infared System (LI-COR, Lincoln, Nebraska) according to manufacturer’s suggestions. Briefly, membranes were blocked in Odyssey blocking buffer and membranes were incubated with 1:200 anti-MMP2 (13595, Santa Cruz), and 1:10,000 anti-beta tubulin (2128, Cell Signaling) overnight in 1% BSA-PBS at 4°C. For detection, secondary antibodies were diluted in 1% BSA-PBS as follows: MMP2 blot with 1:2000 goat anti-mouse IRdye® 800CW, and beta actin with 1:15,000 goat anti-rabbit IRdye® 800CW (LI-COR).

### MT1-MMP ELISA

HPAAF were seeded at a density of 60,000 cells/mL in a 24-well plate in serum media and incubated overnight to allow cell attachment. Media was then changed to serum-free media. Cells were placed in either a standard incubator (normoxia) or in the hypoxia chamber (hypoxia) for 48 hours [Noxygen, Elzach, Germany]. After 48 hours, media was collected and stored at −80⁰C. Media samples were thawed on ice prior to analysis with the ELISA. The Human MMP-14 or MT1-MMP detection was done using an ELISA kit (XpressBio, Frederick, Maryland). Sample incubation, washing, and reagent preparation were conducted according to the kit instructions. Protein concentrations were quantified using absorbance measurements recorded at a wavelength of 450 nm on a Varioskan LUX multimode microplate reader (Thermofisher).

### ATP Assay in cells

HPAAF were seeded at a density of 15,000 cells/well in 96-well plates in serum-free media. After 24 hours, media was changed to media with serum (10% FBS) or media with serum and HIF-1alpha inhibitor (30 µM). Plates were incubated for 48 hours in hypoxia or normoxia. Plates were then allowed to cool at room temperature for 30 minutes in hypoxia or normoxia before using Cell Titer Glo Assay (Promega, Madison, WI). Assay substrate (100 µL) was then added to each well, incubated in dark at room temperature at 10 minutes, and read in luminescence in a spectrophotometer (Infinite M1000 PRO, Tecan).

### Lactate Assay in cells

HPAAF were seeded at a density of 180,000 cells/ml in 24 well plates in serum-free media. After 24 hours, media was changed to media with serum (10% FBS) or media with serum and HIF-1alpha inhibitors (3 µM or 30 µM). Plates were incubated for 48 hours in hypoxia or normoxia. Media was then collected, and lactate assay was performed (Trinity Biotech USA, Jamestown, NY). Briefly, media was diluted 1:10 in PBS and 2 µL of diluted media was added to 200 µL of reagent in each well of a 96 well plate. Plates were incubated for 1 hour and read at 540 nm in a spectrophotometer (Infinite M1000 PRO, Tecan).

### Collagen I ELISA

HPAAF were seeded at a density of 20,000 cells/well in 24 well plates in serum-free media and then treated with 30 µM HIF1alpha inhibitor under normoxic or hypoxic conditions, as described above for the lactate assay. After 48 hours, wells were washed twice with PBS, fixed in 10% formalin for 15 minutes, blocked overnight at 4°C with 3% goat serum in PBS and then incubated in 1:1000 anti-collagen, type I (Clone COL-1, Sigma) in 1% goat serum in PBS for 2 h at 25°C. After two washes with PBS, wells were incubated with 1:1000 goat anti-mouse IgG (H+L) HRP conjugated secondary antibody (Thermo Scientific Pierce, Waltham, MA) in 1% goat serum in PBS, washed twice in PBS, and developed with 1-Step™ Turbo TMB-ELISA (ThermoFisher Scientific) for 5 minutes protected from light. The absorbance was read at 562 nm using a microplate reader (Infinite® M1000 Pro; Tecan, Switzerland).

### Immunofluorescent Detection of α-smooth muscle actin

HPAAFs were seeded at a density of 20,000 cells/well in 24 well plates in serum-free media and then treated with 30 µM HIF1α inhibitor under normoxic or hypoxic conditions, as described above. After 48 hours, wells were washed twice with PBS, fixed in 10% formalin for 15 minutes, blocked overnight at 4°C with 3% goat serum in PBS and then incubated in 1:500 anti-alpha smooth muscle actin (Clone 1A4, Sigma) in 1% goat serum in PBS for 2 h at 25°C. After two washes with PBS, wells were incubated with 1:1000 AlexaFluor 488 goat anti-mouse IgG (ThermoFisher) for 1 hour, counterstained with DAPI (1 µg/mL, ThermoFisher), and then imaged on a fluorescence microscope (Zeiss Axio Observer Z1).

### Mouse model of hypoxic pulmonary hypertension

Male C57BL6/J mice 10 –12 weeks old, with a body weight of 25.5 ±1.6 g were obtained from Jackson Laboratory (Bar Harbor, ME). Mice were exposed to 10 days of normobaric hypoxia. Hypoxia was created in an environmentally controlled chamber in which nitrogen was mixed with room air until an oxygen concentration of 10% was reached; oxygen levels were measured with a sensor in the chamber (Servoflo, Lexington, MA) as previously described (27). Control, normoxia mice were housed in room air. All mice were exposed to a 12-h:12-h light-dark cycle. All procedures were approved by the University of Wisconsin Institutional Animal Care and Use Committee.

### In vivo right ventricular and pulmonary vascular hemodynamic assessment

Surgical preparation, hemodynamic measurements, and analysis were based on established protocols (27–30). Anesthesia was induced with an intraperitoneal injection of urethane solution (1 mg/g body weight) to maintain heart rate. Mice were then intubated and placed on a ventilator (Harvard Apparatus, Holliston, MA). Systemic pressure was measured with a pressure catheter (Millar, Houston, TX) inserted from the common carotid artery. Systemic pressure and heart rate were recorded throughout the procedure. The thoracic cavity was opened, as previously described, and the heart was exposed by removal of anterior rib cage (27, 28, 30). RV pressure-volume loops were obtained as previously described using a 1.2 F admittance catheter inserted through the apex of the heart into the RV. After instrumentation was established and baseline pressure-volume measurements were obtained, the inferior vena cava (IVC) was isolated and intermittently occluded to obtain alterations in venous return for determination of end-systolic and end-diastolic pressure relations. Commercial software (Notocord, Croissy Sur Seine, France) recorded RV pressure and volume waveforms simultaneously, and data were analyzed using a minimum of 10 consecutive cardiac cycles. Cardiac output (CO) was normalized by body weight (BW) to calculate the cardiac index (CI) (27–31).

Pulmonary vascular function was quantified using total pulmonary vascular resistance (TPVR) and pulmonary arterial elastance (E_a_) (30, 32). RV mechanical function was assessed using established parameters including maximum and minimum pressure derivatives (dP/dt_max_, dP/dt_min_), end systolic elastance (E_es_), relaxation factor τ, ejection fraction (RV EF), and chamber compliance as previously described (27, 28, 33). Ventricular–vascular interactions were assessed using E_es_/E_a_ (27, 28).

#### ATP Assay in tissue

ATP in frozen lung tissue was measured colorimetrically using an ATP assay kit (ab83355, Abcam, Cambridge, MA). Roughly 10 mg of lung tissue was dissected, weighed, and rinsed in pre-chilled PBS according to the manufacturer’s protocol. After tissue homogenization, samples were deproteinated and then neutralized using the Deproteinizing Sample Preparation Kit – TCA (ab204708, Abcam, Cambridge, MA). To correct for any sample matrix interference in the sample wells, 1 nmol of ATP was spiked in the sample wells. Duplicate wells were performed for the samples, sample background controls, spiked ATP samples, and ATP standards. ATP concentration was calculated as instructed by manufacturer’s protocol and normalized to wet tissue weight.

### Lactate Assay in tissue

Lactate concentration in frozen lung tissue was measured colorimetrically using a lactate assay kit (73510, Trinity Biotech, Jamestown, NY). Roughly 10 mg of lung tissue was dissected and weighed. Tissue was homogenized in pre-chilled lactate reagent. Duplicate wells were performed for the samples and lactate standards. Lactate concentration was calculated as instructed by the manufacturer’s protocol and normalized to protein concentration.

### Statistics

All data are expressed as mean +/- standard deviation. Student’s t-test or 2-way ANOVA with Tukey’s multiple comparison correction were performed as indicated to determine significance. All experiments were performed at least n=3.

## Results

### Hypoxia drives HPAAF collagen accumulation by reducing turnover

In chronic HPH rodent models, hypoxia causes collagen accumulation in pulmonary arteries (5, 6). Collagen type I is the major fibrillar collagen present in pulmonary arteries in HPH and affects arterial stiffness in the disease. Figure 1A demonstrates that Collagen 1 content increased in cultured HPAAF in response to hypoxia. Additionally, Collagen 1 mRNA levels showed no change in response to hypoxia (Figure 1B), suggesting that the increase in collagen content was not due to an increase in collagen synthesis. Matrix metalloproteinases (MMPs) degrade collagen and play key roles in the development of pulmonary hypertension (34). In HPAAF exposed to hypoxia, MMP activity was significantly reduced (Figure 1C). Specifically, expression of MMP2 was dramatically reduced in hypoxic conditions (Figure 1D). MT1-MMP concentration was also reduced with hypoxia, but was not found to be significantly different from normoxic conditions (Figure 1E). Meanwhile, expression of alpha smooth muscle acting (αSMA), an indicator of differentiation to a myofibroblastic phenotype, was also reduced in hypoxic conditions (Figure 1F). Together, these data indicate that Collagen 1 accumulation in HPAAF in response to hypoxia is driven by a reduction in collagen degradation or turnover rather than an increase in collagen production.

**Figure 1:**
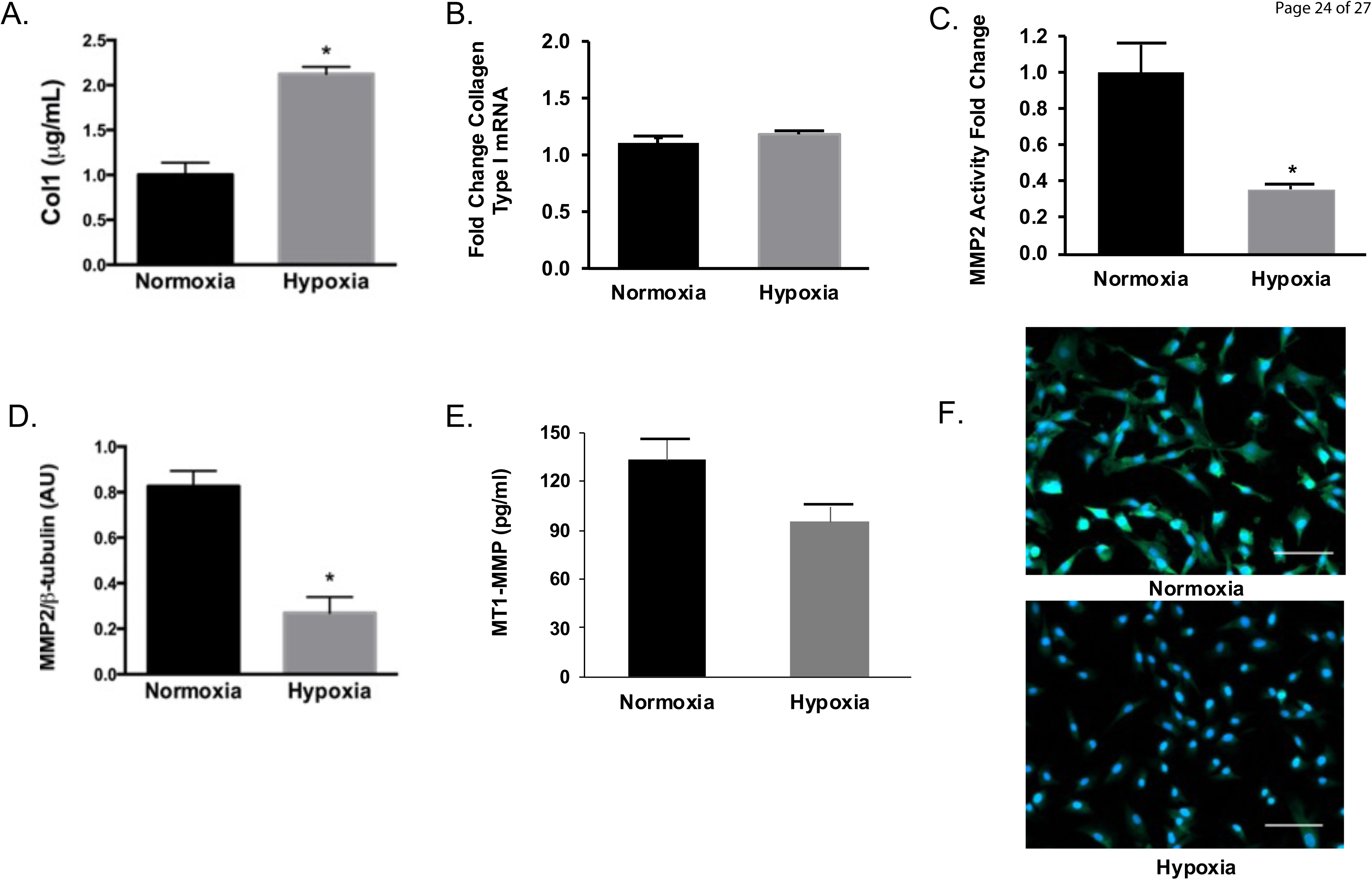
Hypoxia induces a maladaptive PAAF phenotype. A. Collagen I amount determined by ELISA increased in hypoxia, n=3 per group, *p<0.05 vs Normoxia, B. Collagen mRNA expression unchanged in hypoxia, n=4 per group, p=0.728 Hypoxia vs. Normoxia, C. MMP2 activity decreased in hypoxia, n=4 per group, *p<0.05 vs. Normoxia, D. MMP2 expression determined by western blot decreased in hypoxia, n=3, p<0.05 vs Normoxia. E. Concentration of MTP1-MMP decreased in hypoxia, but was not significantly different n=3, p > 0.05. F. Immunofluorescent staining for DAPI (blue) and αSMA (green). Scale bar = 100 μm.

### HPAAF demonstrate altered metabolism and increased HIF-1α expression in response to hypoxia

Metabolic changes in HPAAF exposed to hypoxia were evaluated by examination of lactic acid and ATP concentrations. In response to hypoxia, HPAAFs exhibited a significant increase in lactic acid concentration (Figure 2A) and a reduction in ATP (Figure 2B). In addition to changes in Collagen 1 levels and metabolism, HIF-1α expression was significantly increased in HPAAF in response to hypoxia.

**Figure 2:**
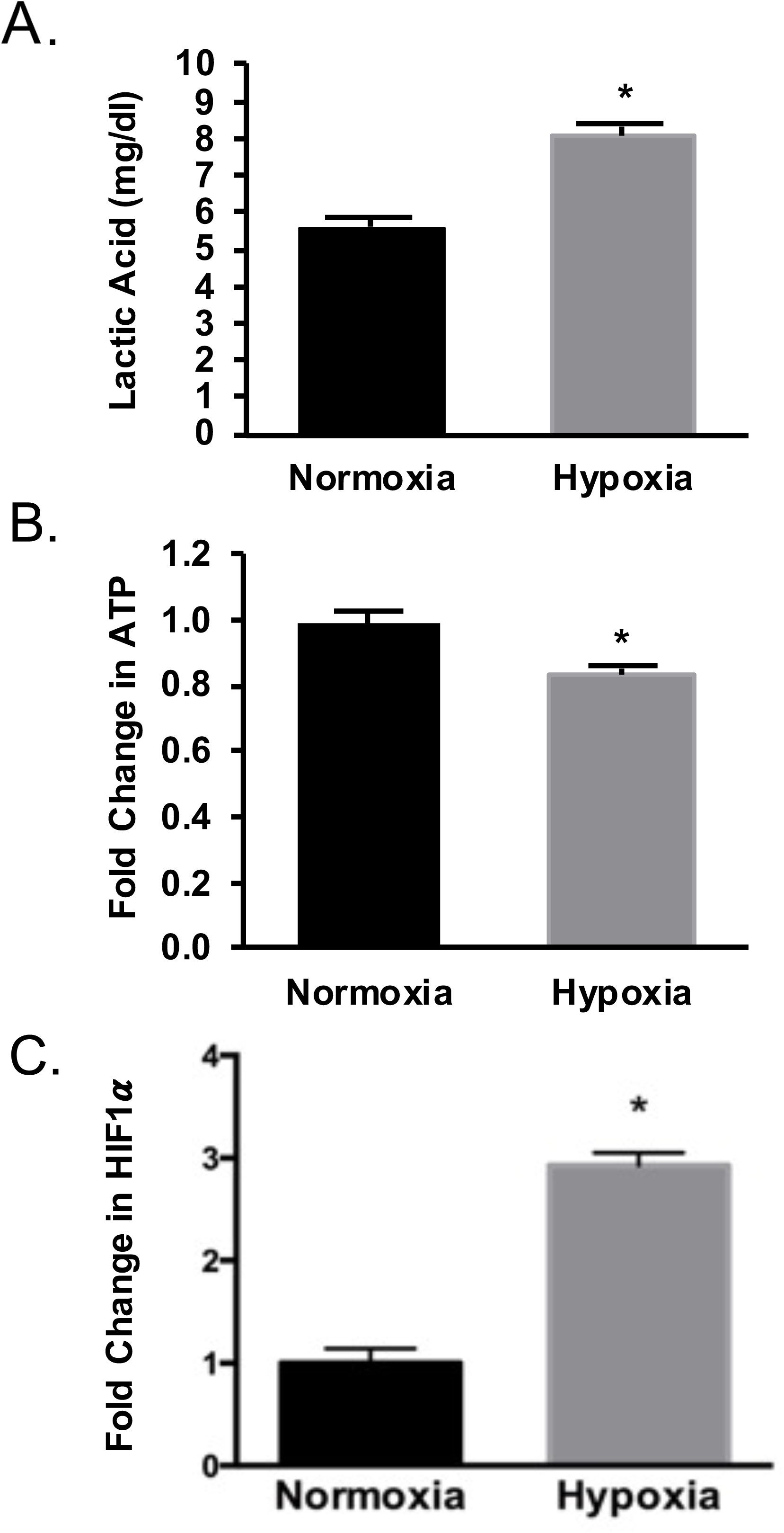
Altered metabolism in PAAF due to hypoxia. A. Lactic acid concentration increase in hypoxia, n=6 per group, *p<0.001 vs. Normoxia, B. Decreased ATP in hypoxia, n=4 per group, p<0.05 vs Normoxia, C. Increased HIF1α expression, determined by ELISA, in hypoxia, n=3, p<0.05.

### HIF-1α inhibition preserves HPAAF phenotype in response to hypoxia exposure

The role of HIF-1α in regulating the changes in Collagen 1 degradation and cellular metabolism in response to hypoxia was investigated using the HIF1-α inhibitor. Cells were treated with 30 µM of inhibitor prior to normoxia or hypoxia exposure. HIF-1α inhibition prevented Collagen 1 accumulation in response to hypoxia (Figure 3A). Consistent with this finding, the hypoxia-induced downregulation of MMP activity and MMP2 expression was also reversed upon inhibition of HIF-1a (Figures 3B&C), and expression of αSMA appeared to increase to levels comparable with normoxia based on examination of immunostained cells (Figure 3D). Similarly, HIF-1α inhibitor treatment preserved lower lactic acid levels in HPAAF exposed to hypoxia (Figure 3E). Furthermore, HIF-1α inhibition preserved ATP levels in human PAAF exposed to hypoxia at levels similar to normoxic HPAAF (Figure 3F). These data demonstrated that the HPAAF metabolism, ATP and lactate production, is preserved in hypoxia by HIF-1α inhibition.

**Figure 3:**
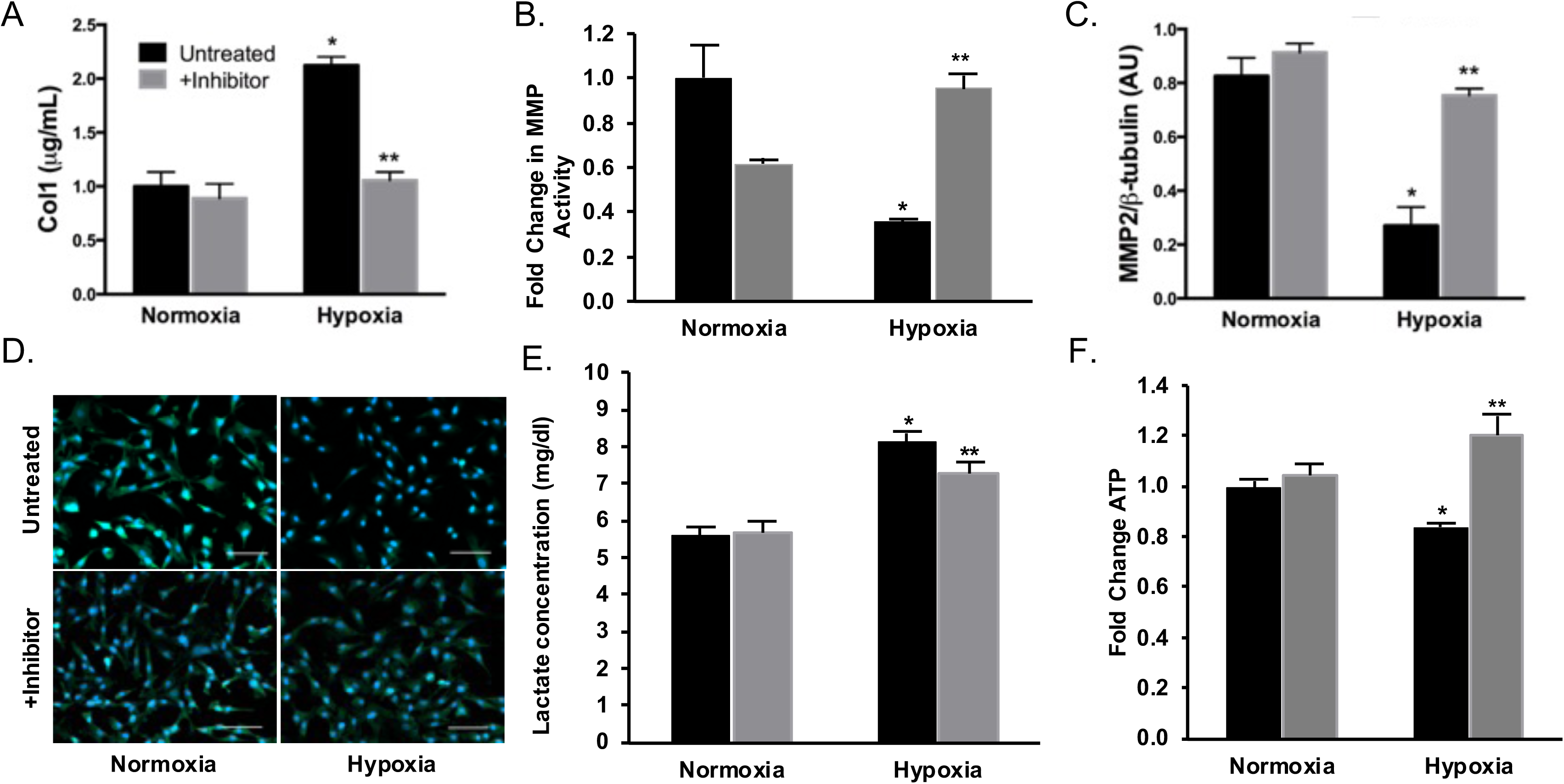
Preservation of HPAAF phenotype with HIF-1α inhibition. A. Col1 mRNA expression in hypoxic HPAAF decreased with HIF-1α inhibition, n=4 per group, *p<0.05 vs normoxia, **p<0.005 vs untreated, B Fold change in MMP activity in hypoxic HPAAF increased with HIF-1α inhibition, n=4 per group, *p<XX vs normoxia, **p<0.05 vs. untreated. C. MMP-2 Expression in hypoxic HPAAF increased with HIF-1α inhibition, n=3 per group, *p<0.05 vs normoxia, **p<0.005 vs untreated. D. Immunostaining for DAPI (blue) and αSMA (green), Scale bar = 100 μm. E. Lactate concentration in hypoxic HPAAF was decreased with HIF-1α inhibition, n=3 per group, *p<0.001 vs. normoxia, **p<0.05 vs. untreated. F. Fold change of ATP in hypoxic HPAAF increased with HIF-1α inhibition, n=4 per group, *p<0.05 vs. normoxia, **p<0.001 vs untreated.

### Altered metabolism in lungs of hypoxic pulmonary hypertension mice

Having demonstrated both changes in Collagen 1 accumulation and metabolism in HPAAF exposed to hypoxia *in vitro*, we next investigated whether similar alterations are observed *in vivo* in a mouse model of HPH. Table 1 demonstrates the development of pulmonary hypertension in mice exposed to hypoxia with a significant increase in RVSP and TPVR compared to control. Table 1 further demonstrates the development on pulmonary vascular mechanical changes with increased pulmonary arterial elastances (E_a_). Several previous studies have demonstrated that collagen accumulation accumulates in the pulmonary arteries in response hypoxia resulting in PA stiffening and increased afterload (5, 6, 8, 35, 36). Following hemodynamic assessment, which confirmed the development of HPH, metabolic changes were evaluated in whole lung homogenates. While lactic acid concentration was preserved in hypoxic lung tissue (Figure 4A), there was a significant reduction in ATP content (Figure 4B), similar to changes in ATP levels seen in HPAAF exposed to hypoxia.

**Figure 4:**
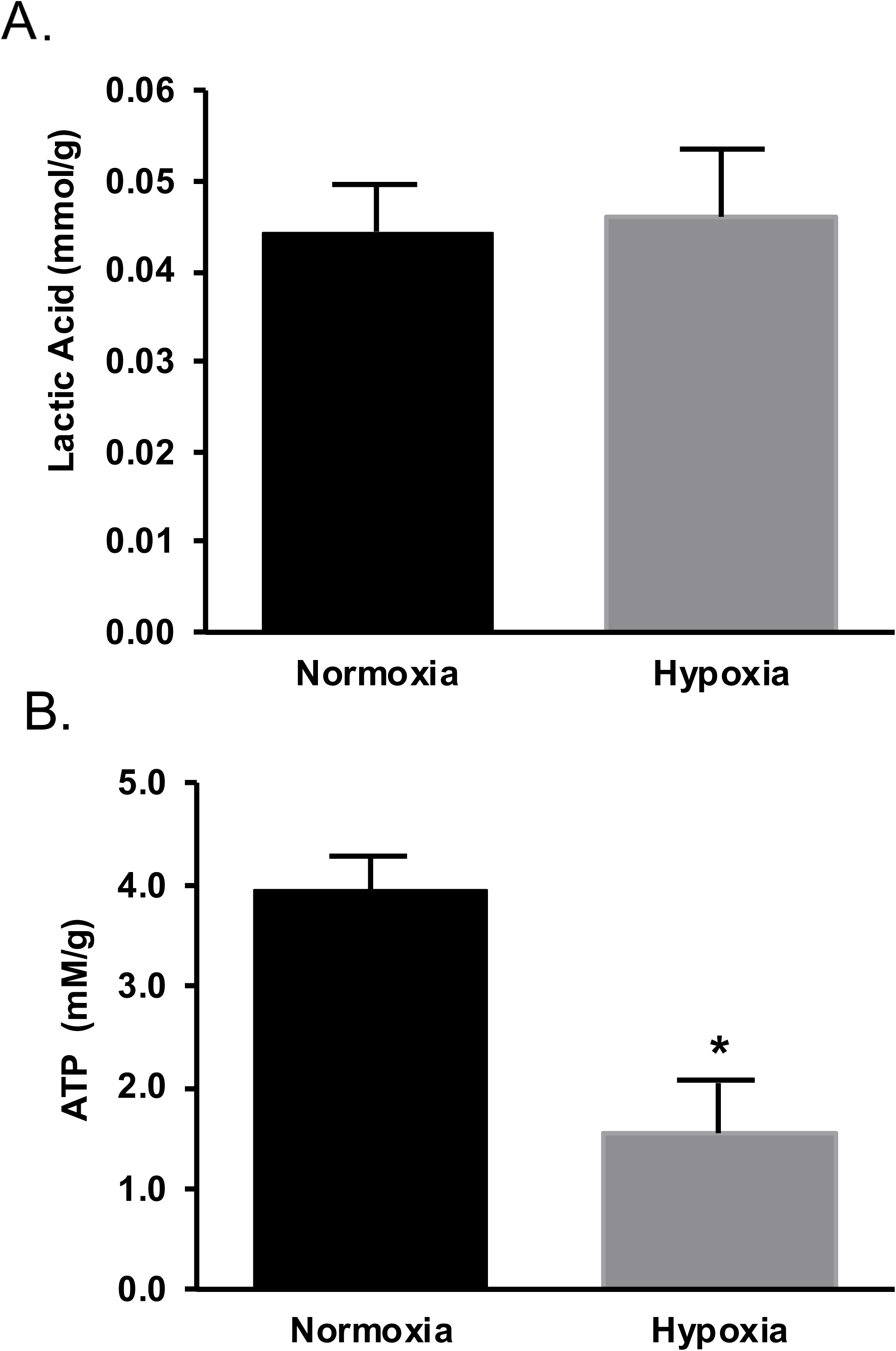
Pulmonary metabolic changes in hypoxic PAH in mice. A. Lactic acid concentration per gram protein in hypoxic PAH mouse lungs, n=5 per group. B. ATP concentration per gram of hypoxic PAH lung tissue, n=5 per group, *p<0.05 vs. Normoxia

**Table 1:**
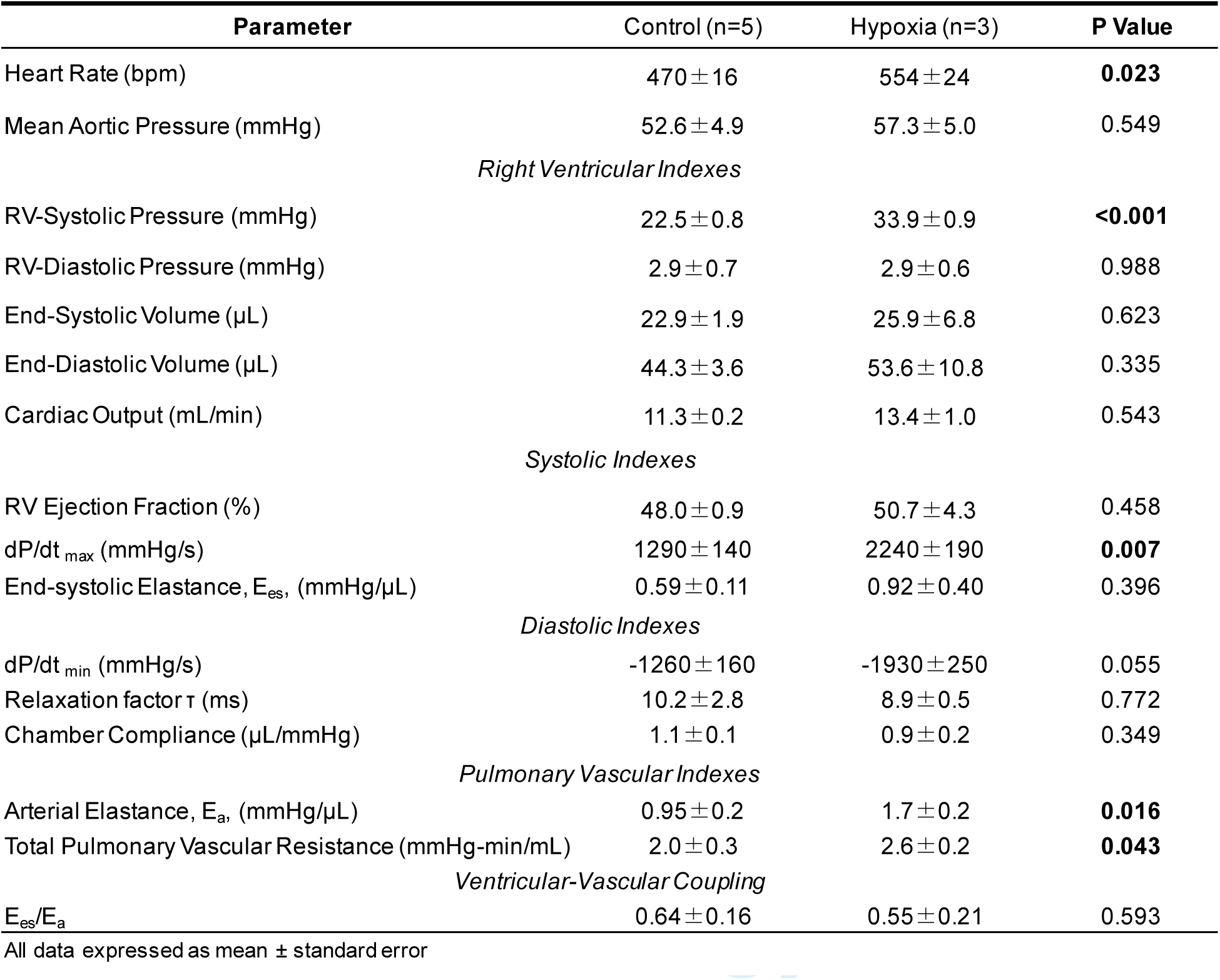
Hemodynamic Analysis of the Right Ventricle and Pulmonary Vasculature.

## Discussion

We have previously demonstrated the importance of collagen accumulation in hypoxic pulmonary hypertension through use of animal models (5). More specifically, we found that increases in collagen amounts in hypoxic pulmonary hypertension lead to increased stiffness of the pulmonary arteries (5). However, the mechanism by which this increase in collagen amount occurs is unclear. This paper is one of the first to specifically elucidate mechanisms of adventitial fibroblast collagen production in hypoxia, along with changes in adventitial fibroblast metabolism.

In this current work, we aimed to elucidate the connection between fibroblast-driven collagen production and degradation and changes in fibroblast metabolism. Our results demonstrate that decrease MMP activity contributes to increased collagen amounts in adventitial fibroblasts in hypoxia while collagen synthesis by the fibroblasts remains unchanged. In addition, our results show that the decrease in MMP activity with hypoxia is dependent on the activity of HIF-1α, part of a heterodimeric transcription factor that is responsible for expression of proteins in hypoxic conditions.

Additionally, we have shown that HIF-1α activation in hypoxia leads to increased lactic acid and decreased ATP production in HPAAF. These findings are consistent with previous studies which have shown that HIF-1α activation increases glycolytic rate, which leads to lactic acid production in various diseases, including cancer (20, 21, 37). Furthermore, ATP has been shown to upregulate MMPs in many contexts (38–40) including stimulating the release of MMP2 from human aortic smooth muscle cells (41). Therefore, it is suggested that the decrease in ATP may be linked to the decrease in overall MMP2 activity. Furthermore, the concentration of MT1-MMP, while not significantly lower, was decreased in HPAAF exposed to hypoxia. MT1-MMP is responsible for the activation of MMP2 and has been found to regulate ATP levels through stimulation of the HIF-1α pathway in other disease states (42) suggesting a potential hypoxia-driven signaling mechanism for the decreased MMP2 activity and expression that merits further investigation.

Hypoxia has been shown to alter metabolism of pulmonary arterial cells in hypoxic pulmonary hypertension(4). Both pulmonary arterial adventitial fibroblasts and macrophages subjected to hypoxia demonstrate increased glycolysis compared to normoxia (4). The increase in aerobic glycolysis in hypoxia has also been demonstrated for endothelial cells and smooth muscle cells in the pulmonary artery (4). The increase in aerobic glycolysis agrees with our results showing increases in lactate media concentration in hypoxia, which suggests that hypoxia increases aerobic glycolysis in HPAAF. Hypoxia has also been shown to alter mitochondrial function in pulmonary arterial fibroblasts (43). Specifically, PAAF from neonatal calves subjected to chronic hypoxia had decreased ATP synthesis and mitochondrial complex I activity compared to PAAF from neonatal normoxic calves (43), which agrees with our findings that hypoxia decreased ATP cellular content in HPAAF and suggests that hypoxia decreases mitochondrial function. Our findings are consistent with this previous work. Importantly, our paper is the first to show that these changes in ATP cellular content in hypoxia are mediated through expression of HIF-1α in HPAAF.

Previous studies have shown that hypoxic activation of HIF-1α leads to changes in ECM remodeling, which is in agreement with our results that HIF-1α expression in hypoxia leads to increased collagen I accumulation by HPAAF. Specifically, hypoxia increases the expression of HIF-1α, which leads to increases in ECM deposition and stiffness in human foreskin and breast cancer fibroblasts. The pathway was shown to involve expression of intracellular prolyl and lysyl hydroxylation that stabilizes procollagen molecules, specifically P4HA1, P4HA2, and PLOD2 (14). Additionally, another group demonstrated that in hypoxia, HIF-1α mediates the expression of lysyl oxidase (LOX), an enzyme involved in collagen remodeling. Loss of LOX lead to a decrease in lung cancer aggressiveness and progression(44).

MMPs also are dysregulated in hypoxia-induced pulmonary hypertension (34). Increased MMP expression or activity has been associated with normoxic recovery from hypoxia-induced pulmonary hypertension (45, 46), which supports our finding that hypoxia increases MMP2 activity in HPAAF. In addition, a study by George et al. that demonstrates attenuation of PAH remodeling through transgenic expression of MMP-1 also supports our findings (47).

This study has several key strengths. Importantly, all human cells were used for *in vitro* studies in order to maintain the highest fidelity possible to the human disease states studied. Furthermore, conclusions drawn from *in vitro* studies were confirmed with *in vivo* studies in a small animal model. There are also important limitations in this work. We show strong evidence that collagen accumulation by HPAAF in hypoxia is driven by HIF-1α mediated inhibition of MMP2 expression and activity. However, further investigation into the exact mechanisms by which HIF-1α regulates MMP2 was beyond the scope of this current work and should be a focus of future studies. Additionally, it is likely that accumulation of collagen in HPAAF in hypoxia occurs along multiple pathways and mechanisms, many of which, including inhibition of other MMPs and regulation of TIMPs, are not explored in this current work. *In vivo* HPH in mice provides an important confirmation of many of the *in vitro* findings. However, PAAF were not able to be isolated from hypoxic mouse tissue, therefore, while the same metabolic and fibrotic changes are observed *in vivo* as *in vitro*, we cannot definitely demonstrate what portion of the *in vivo* changes are driven by changes in PAAF. Additionally, further studies should explore the impact of HIF-1α inhibition on development of HPH *in vivo* in order to better understand its potential as a therapeutic target.

In summary, we have shown that hypoxia leads to collagen accumulation in HPAAF by decreasing MMP activity and expression in an HIF-1α dependent manner, possibly via MT1-MMP related signaling pathways. Additionally, we have shown that hypoxia increases lactic acid and decreases ATP content through HIF-1α activation. We further showed that this increase in ATP is observed in an *in vivo* model of HPH. Since ATP has been shown to increase MMP activity, this suggests that the decrease in ATP content could potentially be responsible for decreased MMP2 activity, leading to collagen accumulation mediated by HPAAF. This work presents a potential mechanism by which hypoxia induces collagen accumulation in the pulmonary artery adventitia in hypoxic pulmonary hypertension, which contributes to arterial stiffening, increased RV afterload and ultimately RV failure.

## Acknowledgements

This work was supported, in part, by a Thoracic Surgery Foundation Nina Starr Braunwald Research Fellowship (to J.L.P.), the National Institutes of Health (R01-HL086939 to N.C.C, and R01-HL141181 to K.S.M). The authors would like to thank Nikhita Chawla for her assistance in conducting cell culture experiments.

## Author Contributions

J.L.P., C.A.C., K.S.M., and N.C.C. conceived and designed research; J.L.P, C.A.C., L.R.C., N.P., T.C.C., N.C., K.A.B, D.M.T., T.M., and T.A.H conducted experiments for results and prepared figures; J.L.P, C.A.C., L.R.C., K.S.M, T.M., and N.C.C. drafted and revised manuscript; N.C.C. approved manuscript.

